# Neo-natal castration leads to subtle differences in porcine anterior cruciate ligament morphology and function in adolescence

**DOI:** 10.1101/2023.01.24.524954

**Authors:** Jacob D. Thompson, Danielle Howe, Emily H. Griffith, Matthew B. Fisher

**Author notes:** Corresponding Author Contact: Address: 4130 Engineering Building III, 911 Oval Drive, CB 7115, Raleigh, NC, 27695. **Author Contribution** Jacob D. Thompson contributed to the development of research design, acquisition, analysis, and interpretation of data, and drafting of the manuscript. Danielle Howe contributed to the development of research design, acquisition of data, and revision of the manuscript. Emily H. Griffith contributed to the interpretation of data and revision of the manuscript. Matthew B. Fisher contributed to the development of research design, acquisition, analysis, and interpretation of data, and revision of the manuscript. All authors have read and approved the final submitted manuscript.

## Abstract

Female adolescent athletes are at a higher risk of tearing their anterior cruciate ligament (ACL) than male counterparts. While most work related to hormones has focused on the effects of estrogen to understand the increased risk of ACL injury, there are other understudied factors, including testosterone. The purpose of this study was to determine how surgical castration in the male porcine model influences ACL size and function across skeletal growth. Thirty-six male Yorkshire crossbreed pigs were raised to 3 (juvenile), 4.5 (early adolescent), and 6 months (adolescent) of age. Animals were either castrated (barrows) within 1-2 weeks after birth or were left intact (boars). Post-euthanasia, joint and ACL size were assessed via MRI, and biomechanics were assessed via a robotic testing system. Joint size increased throughout age, yet barrows had smaller joints than boars (p<0.001 for all measures). ACL cross-sectional area (CSA), length, volume, and stiffness increased with age (p<0.0001), as did ACL anteromedial (AM) bundle percent contribution to resisting loads (p=0.012). Boar ACL, AM bundle, and PL bundle volumes were 19% (p=0.002), 25% (p=0.003), and 15% (p=0.04) larger than barrows across ages. However, CSA, stiffness, and bundle contribution were similar between boars and barrows (p>0.05). The barrows had smaller temporal increases in AM bundle percent function than boars, but these data were highly variable. Thus, early and sustained loss in testosterone leads to subtle differences in ACL morphology, but may not influence measures associated with increased injury risk, such as CSA or bundle forces in response to applied loads.

## Introduction

There are an ever-increasing number of pediatric athletes participating in middle and high school sports^1^, accompanied by an increase in pediatric anterior cruciate ligament (ACL) injuries^2–4^. Furthermore, during adolescence there is a difference in the rate of ACL injuries between males and females, with females being 1.4 times more likely to tear their ACL overall^5^. In specific sports, they are over 3 times more likely to tear their ACL, like basketball^6–8^ or soccer^9^. Several groups attribute this difference to aspects of sexual maturity^10,11^ or gendered expectations in training, sport selection, and coaching^12,13^. A large focus has been on the onset of spikes and cycles of sex hormones^14–17^. The female sex hormones estradiol and progesterone have gathered the most attention with some groups showing associations between peaks of estradiol and progesterone to knee joint laxity and time of ACL injury^13,14,16,18^. However, strong evidence for a direct correlation between estradiol levels and injury risk does not yet exist^16,17^. While a major focus has been on uncovering theorized catabolic effects of female sex hormones to understand the increased risk of ACL injury, an understudied factor is the influence of testosterone, the primary male sex hormone, on developing male ACLs.

It is known that there are androgen receptors present within the human male ACL^19^, suggesting that the ACL is an androgen-responsive tissue. In humans, males who tore their ACLs have higher levels of testosterone, indicating that testosterone may play some direct or indirect role in increasing ACL injury risk^20^. To study testosterone in animal models, surgical castration has been performed to drastically decrease total serum testosterone by over 95%^21,22^. For example, adult rats castrated at 12 weeks of age (post-puberty) had a 25-fold lower testosterone at 5 weeks post-castration than age-matched controls. Although the ACLs in the castrated rat group had a similar size, they had 15% lower load-to-failure values and 18% lower ultimate stresses than intact males^22^. This study provides some evidence that testosterone may play a protective role in ligament function in adults, especially failure properties of the ACL^22^. Thus, it may be possible that both high and low levels of testosterone may impact the ACL, yet the evidence is still sparse, particularly during skeletal growth.

Large animal models provide a useful tool to study the ACL. Our group, along with others, have used a porcine model to understand structural and functional changes of the ACL throughout skeletal growth, including progression through adolescence^23–28^. We have shown similar size and functional changes within the human and porcine ACL, making this a good model to study changes during adolescence^23,29^. The size of the porcine ACL diverges in males and females around the point of sexual maturity, mainly within the two distinct bundles of the ACL, the anteromedial (AM) and posterolateral (PL) bundles^30^. The AM bundle in males and females increase in size throughout adolescence, yet the PL bundle size plateaus near the onset of puberty. Functionally, there is also a shift in bundle function when resisting anterior loads, where the AM bundle becomes the primary stabilizer over the PL bundle in both males and females^30^. The time of these changes is sex-dependent, with the plateau in PL bundle size occurring earlier and functional changes occurring later in females. These sex specific changes, and their timing around adolescence, motivate the examination of the role of sex hormones as a potential cause of these changes. It is still unclear, though, which sex hormones drive changes in males and females. Therefore, the objective of this study was to determine how neo-natal castration influences the structure and function of the male porcine ACL during youth and early adolescence.

## Methods

### Specimen Collection and Structural Imaging

Thirty-six male Yorkshire crossbreed pigs were raised to 3 months (juvenile), 4.5 months (pre-puberty), or 6 months (early puberty). Six of each age group were left sexually intact (boars), while the other six were castrated (barrows) at an early age (n=6 animals/age/treatment). The boar data presented in this study has been re-analyzed from a previous study^30^ to be compared directly with barrow data. All animals were bred at the North Carolina State University Swine Education Unit, and all experimental protocols were approved by the NC State University IACUC. Following euthanasia, hind limbs were removed, and the stifle (knee) joints were stored at −20°C in saline-soaked gauze.

To determine structural data, stifle joints were thawed and imaged using a 7.0-T Siemens Magnetom MRI scanner (Siemens Healthineers, Erlangen, Germany) using a double-echo steady-state sequence (DESS, flip angle: 25°, TR: 17ms, TE: 6s, voxel size: 0.42 × 0.42 × 0.4mm). Joint size data, including bicondylar width, tibial CSA, anterior-posterior (AP) tibial width, as well as ACL, AM and PL bundle CSAs, orientations, lengths, and volumes were recorded after segmentation using a commercial software (Simpleware ScanIP, Synopsys, Mountain View, CA, USA) and processing using MATLAB (MathWorks, Natick, MA, USA) scripts. Morphological ACL characteristics, including CSA, length, and volume, were normalized to bicondylar width to account for overall joint size.

### Biomechanical Data Collection and Processing

Stifle joints were thawed, trimmed, and the femur and tibia mounted in custom molds using a fiberglass reinforced epoxy (Everglass, Evercoat, Cincinnati, OH, USA) to prepare for robotic biomechanical testing. Stifle joints were then mounted to a robotic testing system (KR300 R2500, KRC4, Kuka, Shelby Charter Township, MI, USA) with a universal force-moment sensor (Omega160 IP65, ATI Industrial Automatic, Apex, NC, USA), following previously described methods^30^. All systems were controlled through the simVITRO software knee module (Cleveland Clinic, Cleveland, OH, USA).

During each test, the femur was fixed to the testing platform and the tibia was mounted to the robotic system. Anatomical points on each leg were recorded using a 3D digitizer (G2X MicroScribe, Amherst, VA, USA) to establish an anatomic coordinate system. The joints were then flexed from 40° (approximately full extension in pigs) to 90° while minimizing joint forces to establish a passive path during the test. AP forces were then applied to the joints at 60° of flexion, while minimizing forces in other directions and recording kinematic data. Maximum loads were scaled across age (40 N, 80 N, and 100 N for 3-month, 4.5-month, and 6-month, respectively) based on the femoral cortical CSA. The recorded kinematics were then repeated and the forces were measured. Joint tissues were then removed sequentially (capsule, AM bundle, then PL bundle), and the intact kinematics were repeated to determine the force contribution of each tissue through the principle of superposition^31,32^. After the AM bundle was transected, the same loads were applied to the joint while new kinematics were recorded to fully engage the remaining PL bundle.

Force-translation curves were then plotted across AP loading. The engagement point of the ACL was determined by the point of inflection in the force-translation curves, which separated the toe region and the linear region in anterior loading^33,34^. The toe region was fit using an exponential curve, and the higher loads above the inflection point were fit to a line. *In situ* joint slack and stiffnesses were then calculated using Matlab scripts, as previously described^25,33^ (Figure S1). The stiffness values for the ACL and AM bundle were calculated under the intact kinematics, where the PL bundle stiffness was calculated under the AM deficient state to ensure PL engagement. AM and PL contribution percentages to resisting anterior loading were calculated by dividing the anterior force for each bundle by the total ACL contribution to anterior loading and multiplying by 100. The difference in AM bundle contribution and PL bundle contribution were plotted to directly compare the boars and barrows.

### Statistical Analysis

Overall effects of age and surgical castration on bicondylar width, CSA, length, volume, stiffness, slack, and difference in bundle percent contribution were assessed using two-way ANOVAs for each variable. No animals were excluded from this work. Multiple comparisons made between ages and boars and barrows at each age were run and the Type I error rate was controlled using Holm-Sidak’s post hoc method. The difference in AM and PL bundle percent contribution was analyzed using one sample t-tests comparing the means to zero. Overall significance was set to α=0.05 for all tests.

## Results

### Joint Size Decreases following Surgical Castration

From MRI scans, joint bicondylar width, tibial plateau CSA, and AP tibial width were recorded as metrics of joint size between the boars and barrows (Fig 1, Table S1). Across all three measurements of joint size, age had significant effects on overall knee joint size (p<0.0001 for all measures). Specifically, bicondylar width increased by 20% and 21% from 3-months to 6-months for the boars and the barrows, respectively (Fig 1A,D). Tibial plateau CSA increased by 64% and 63% from 3-months to 6-months for boars and barrows, respectively (Fig 1B,E). Similar changes were seen for AP tibial width, increasing by 25% and 32% for boars and barrows between the 3-month and 6-month age groups (Fig 1C,F). Surgical castration had significant effects on joint size (bicondylar width: p=0.001, tibial plateau CSA: p=0.0007, AP tibial width: p<0.0001). Barrow bicondylar width was 8% and 7% smaller at 3-months (adj p=0.03) and 6-months (adj p=0.02) but was not statistically significantly smaller at 4.5 months (adj p=0.40). There were statistically significant (adj p=0.003) differences at 6 months in tibial plateau CSA between boars and barrows with the barrow plateau CSA being 13% smaller on average. All barrow AP tibial widths at each age group were statistically significantly smaller than the boars (3-month: 10% smaller, adj p=0.002; 4.5-month: 5% smaller, adj p=0.05; 6-month: 6% smaller, adj p=0.02).

**Figure 1.**
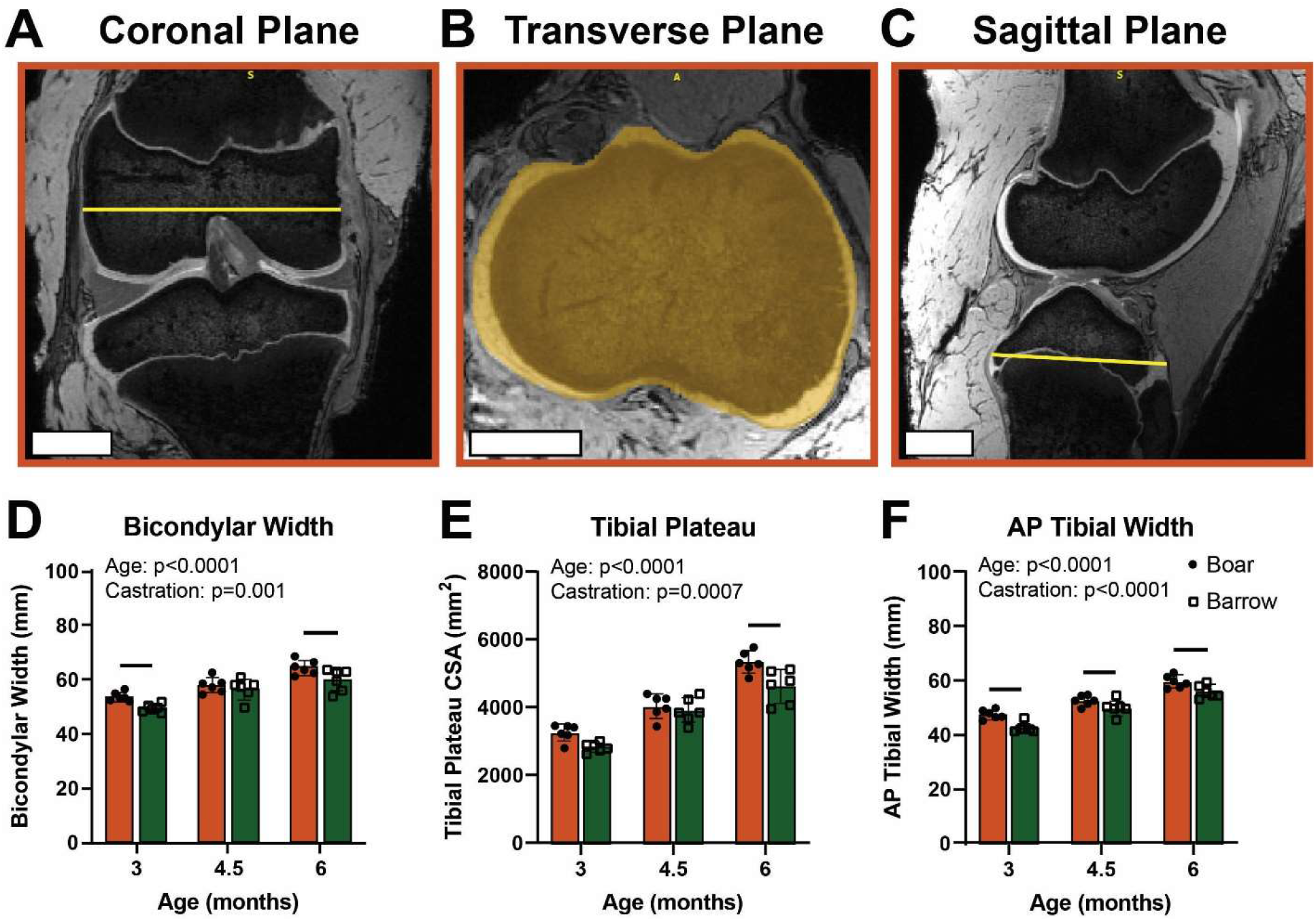
Barrows have smaller joints than boars throughout growth. Joint size measurements taken from the (A) coronal, (B) transverse, and (C) sagittal planes from the MRI scans. Quantitative comparison of (D) bicondylar width, (E) tibial plateau CSA, and (F) AP tibial width measurements between boars and barrows across three ages tested, where each point represents independent specimens. Data presented as mean ± 95% confidence interval with main effects from two-way ANOVA shown in graph corner. Bars indicate p<0.05 between boars and barrows within ages.

### ACL Structure and Morphology Between Castrated and Intact Animals

In addition to overall joint size, the morphology of the ACL and its corresponding AM and PL bundles were recorded. CSAs, lengths, and volumes for the ACL and the bundles were segmented from the MRI scans. For both boars and barrows, there were significant age effects seen for ACL, AM, and PL CSAs (p<0.0001 for all) (Fig 2A, Table S2). When comparing boars and barrows, there was a minor, but not statistically significant, effect of castration on ACL and AM CSA (p=0.12, p=0.20), with boars having slightly higher CSA values at 4.5 and 6-months (12% larger for both ages). At 3 months, boars and barrows had similarly-sized ACLs, but barrows had slightly larger AM bundles (3-months: 3% larger). There was no significant effect of castration on raw PL bundle CSA size (p=0.66). ACL and bundle orientation were also recorded in the sagittal plane, and the orientation of the ACL and the bundles increased across age but were not statistically significant between boars and barrows (Fig S2). All CSA values were normalized to the bicondylar width of the joint to account for overall joint size (Fig 2B). After normalizing, the effect of age persisted for the ACL, AM, and PL CSAs (p<0.0001 for all), but the lack of substantial effect due to castration stayed the same (p=0.81, p=0.70, p=0.27).

**Figure 2.**
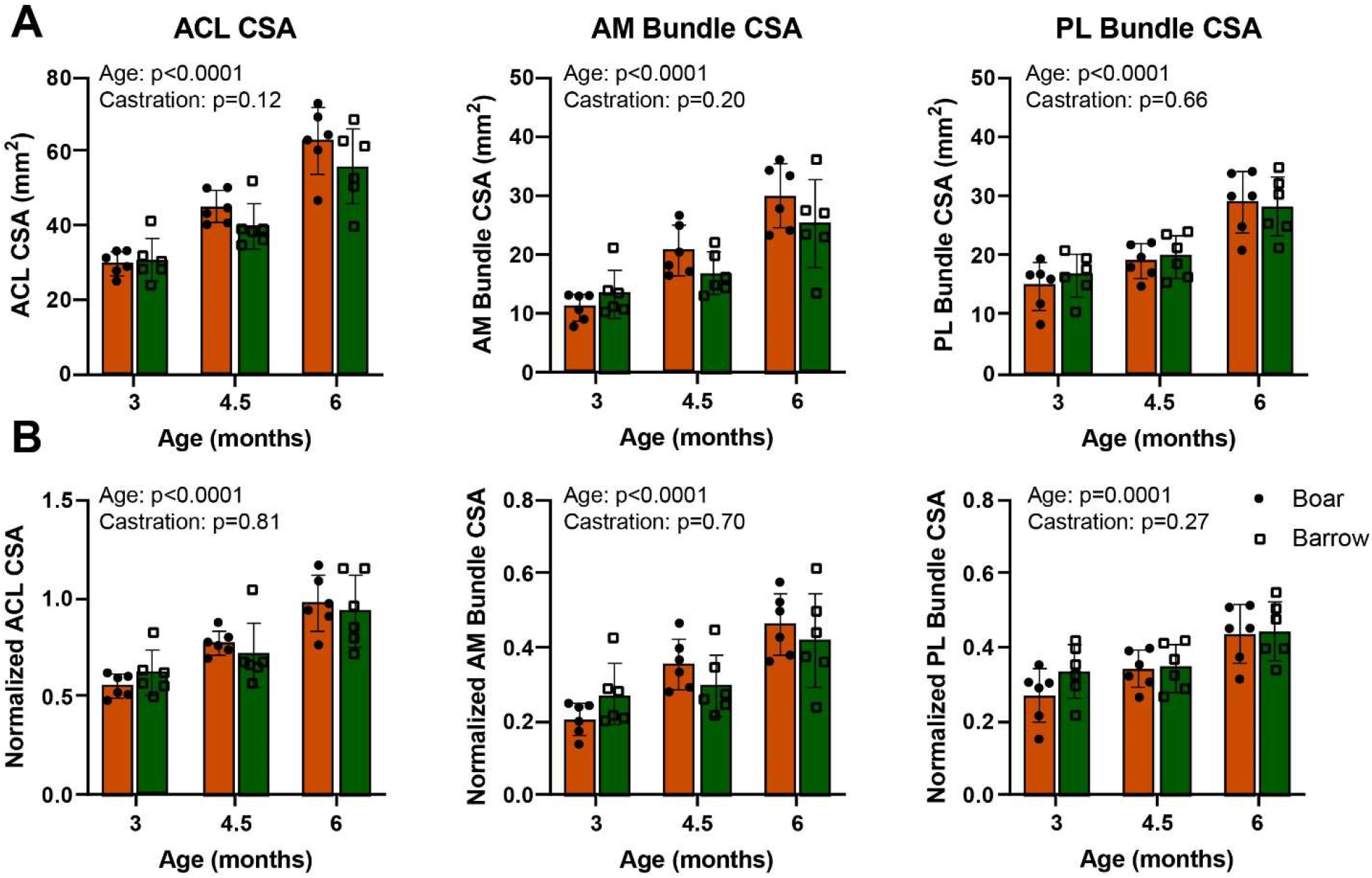
Boar and barrow ACL and bundle CSA increased throughout skeletal growth and adolescence in similar ways. (A) CSAs of the ACL, AM bundle, and PL bundle increased steadily across adolescence, and increases were similar between boars and barrows. (B) Normalizing by bicondylar width to account for overall joint size diminished any minor differences between boars and barrows. Data presented as mean ± 95% confidence interval with main effects from two-way ANOVA shown in graph corner.

Age had similar effects on ACL and bundle lengths, steadily increasing from 3 months to 6 months (p<0.0001 for all) (Fig 3A, Table S3). There was also an effect of castration on the ACL length (p=0.0018) and PL bundle length (p=0.0002), but not the AM bundle length. The boars had on average 8% longer ACLs across all age groups tested. The boars also had longer PL bundles at each age (3-month: 17% greater, adj p=0.005; 4.5-month: 10% greater, adj p=0.07; 6-month: 6% greater, adj p=0.12). After normalizing to joint size, the statistically significant difference in ACL lengths across ages and between boars and barrows disappeared (Fig 3B). The age effect also disappeared for the AM bundle and PL bundle lengths. However, when comparing normalized boar and barrow ACL and bundle lengths, there was still a statistically significant difference in the PL bundle length (p=0.012), with barrows having a statistically significant shorter PL bundle at 4.5 months (11% shorter, adj p=0.02).

**Figure 3.**
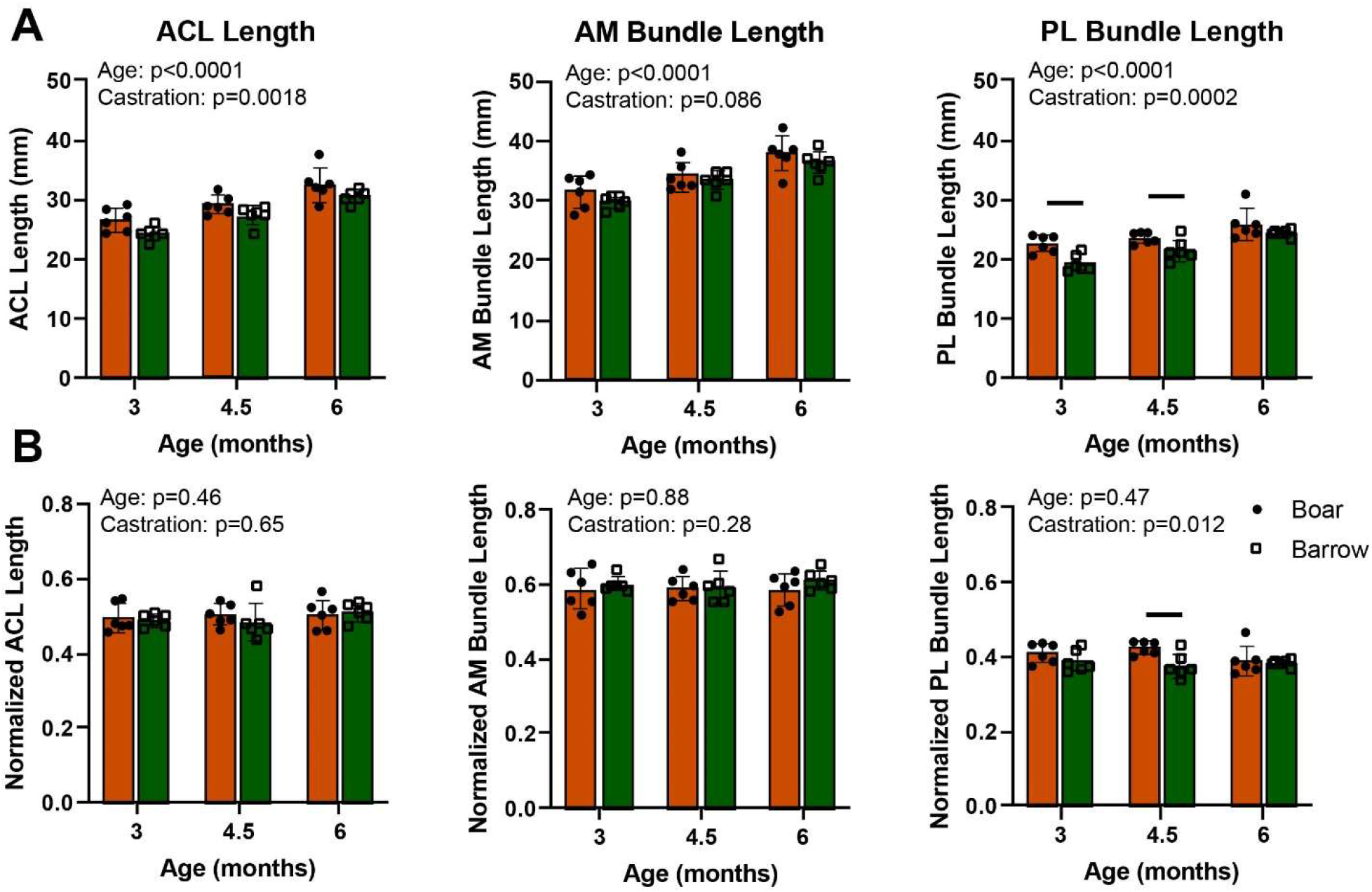
Boar and barrow ACL and bundle lengths all increased throughout skeletal growth and adolescence but had differing lengths. (A) Lengths of the ACL, AM bundle, and PL bundle increased steadily across adolescence, but barrows had shorter ACLs and PL bundles than boars. (B) Normalizing by bicondylar width to account for overall joint size diminished the differences in ACL length, but not PL bundle length, between boars and barrows. Data presented as mean ± 95% confidence interval with main effects from two-way ANOVA shown in graph corner. Bars indicate p<0.05 between boars and barrows within ages.

While ACL and bundle CSA and length showed some minor differences between boars and barrows across the ages tested, the ACL and bundle volumes showed more substantial changes between boars and barrows. A similar age effect was observed in ACL, AM bundle, and PL bundle volumes (p<0.0001 for all), which all increased during growth (Fig 4A, Table S4). Castration also had a statistically significant impact on ACL (p=0.002), AM bundle (p=0.003), and PL bundle (p=0.04) volume. Comparing boars and barrows across ages, the boars had on average 19% larger ACLs, 25% larger AM bundles, and 15% larger PL bundles than the barrows. The age effect is still seen after normalizing to the bicondylar width of the joints, but the effects of castration are no longer statistically significant for the ACL (p=0.06) and PL bundle (p=0.11), but persists in the AM bundle (p=0.04) (Fig 4B).

**Figure 4.**
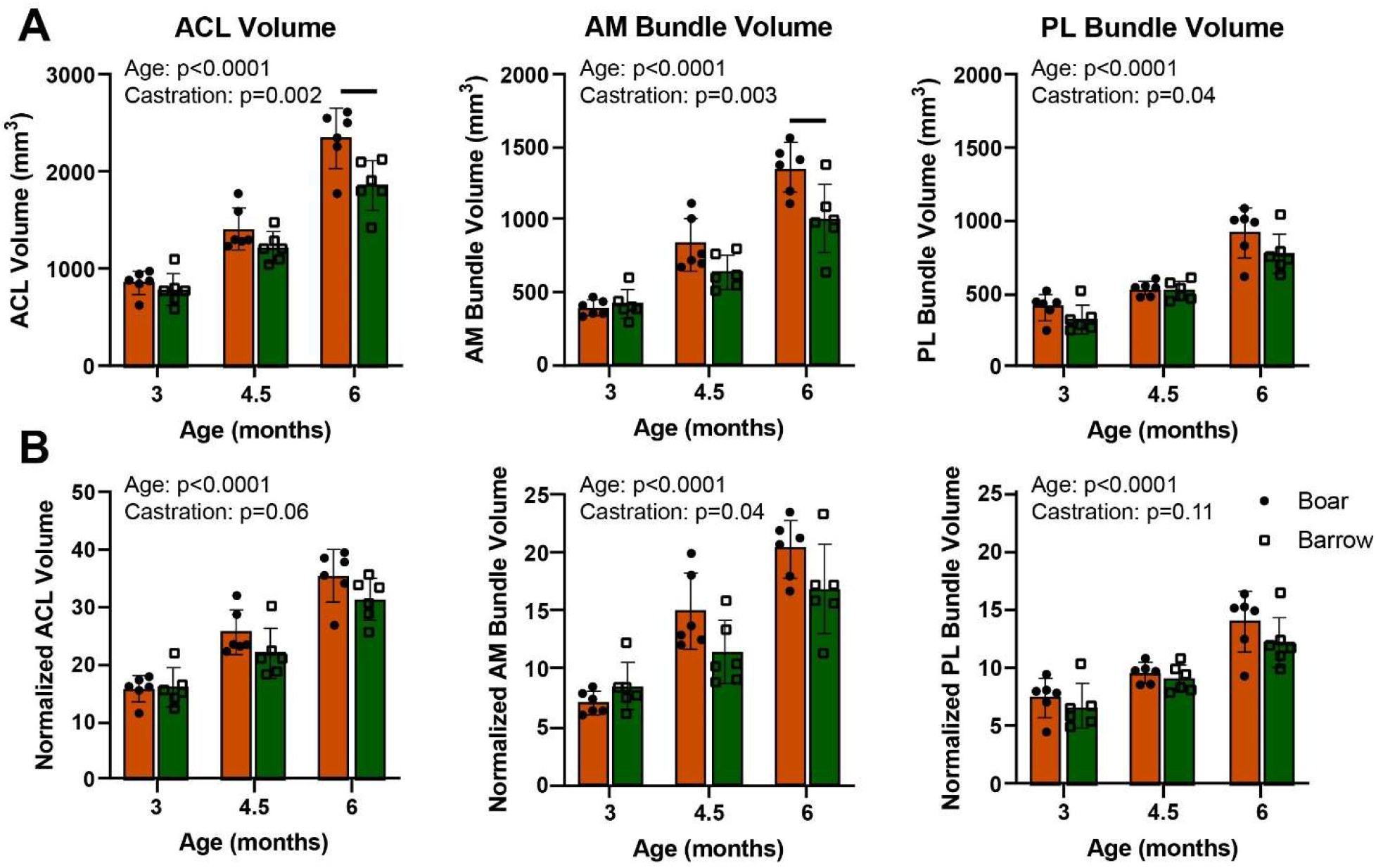
Boar and barrow ACL and bundle volumes all increased throughout skeletal growth and adolescence with barrows having smaller ACL and AM bundle. (A) Volumes of the ACL, AM bundle, and PL bundle increased steadily across adolescence, but barrows had smaller ACLs and AM bundles, but not PL bundles than boars. (B) Normalizing by bicondylar width to account for overall joint size diminished the differences in ACL and AM bundle volume between boars and barrows. Data presented as mean ± 95% confidence interval with main effects from two-way ANOVA shown in graph corner. Bars indicate p<0.05 between boars and barrows within ages.

### Minor Differences in ACL Mechanics in Castrated Animals

Using a 6-degree of freedom robotic testing system, force translation curves showed similar trends in both boars and barrows, where the slopes increased with age, indicating an increase in overall joint stiffness (p<0.0001) (Fig 5A-B). There were no statistically significant differences in overall joint stiffness, however, between boars and barrows (p=0.10) (Fig 5C). Joint slack was calculated as the distance between engagement points of the ACL and the PCL. Both age (p=0.03) and castration (p=0.009) had overall significant effects on joint slack (Fig 5D). The barrows, in general, had shorter slack lengths than the boars, and slack increased throughout growth. Age did not seem to have an effect on the boar group, however, with slacks remaining somewhat similar across ages. Normalized slack values showed no statistically significant effects of age (p=0.40) or castration (p=0.17) (Table S5). Similar changes were seen in the joint laxity, reported as APTT (Fig 5E, Table S6). Both age (p=0.0008) and castration (p=0.008) had pronounced impacts on APTT, with barrows having lower APTT values than the boars, specifically 19% shorter APTT at 3 months (p=0.03). Both groups had increased APTT values across the age groups tested. However, once normalizing to joint size, these age and castration effects disappeared (Fig 5F).

**Figure 5.**
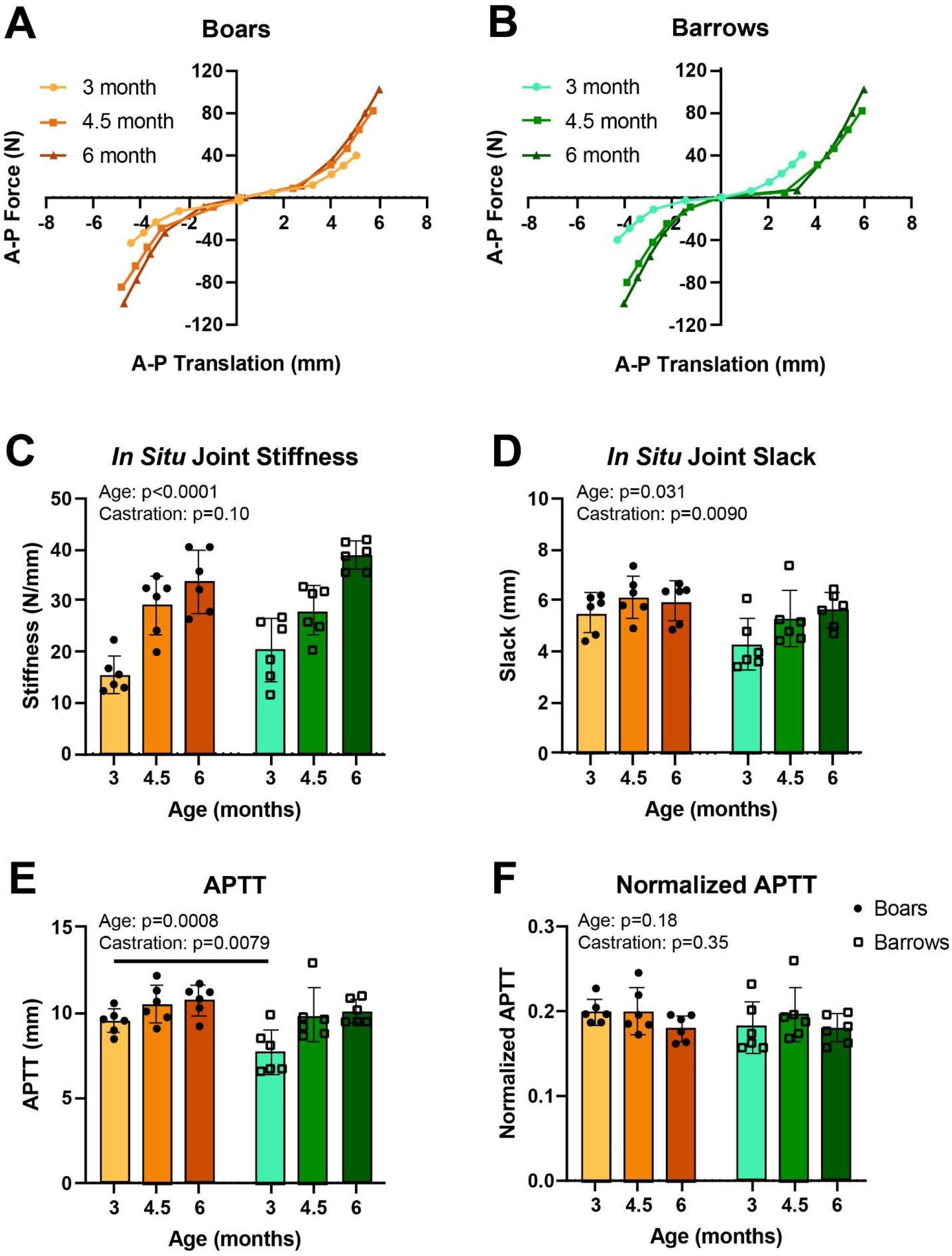
*In situ* joint stiffness, slack, and APTT all change throughout skeletal growth with barrows having less slack and raw laxity than boars. Force translation curves are similar between (A) boars and (B) barrows, with increasing slopes, indicating (C) more stiff joints throughout skeletal growth. (D) *In situ* joint slack slowly increases across age in boars and barrows, but more prominently in the barrow group. (E) Castration and age had significant impacts on joint APTT yet had (F) no significant impact after normalizing to joint size. Force translation data presented as mean forces and mean translations. All other data presented as mean ± 95% confidence interval with main effects from two-way ANOVA shown in graph corner. Bars indicate p<0.05 between boars and barrows within ages.

Similar effects were seen in terms of ACL and bundle function. Stiffness increased with age (ACL: p<0.0001, AM bundle: p<0.0001, PL bundle: p=0.003), but did not change due to castration (ACL: p=0.11, AM bundle: p=0.76, PL bundle: p=0.15) (Fig 6A-C, Table S7). ACL stiffness values for the boars increased by 111% from 3 to 6 months, and the barrow stiffness values increased by 91%. The ACL was the primary contributor to resisting anterior drawer in both the boars and barrows, but the two bundles of the ACL had distinct contributions to resisting load that varied across age (Fig 6D, Table S8). At 3-months, the AM bundle contributed slightly more to resisting anterior drawer than the PL bundle for both boars and barrows. From 3-months to 4.5-months, the AM bundle in the boars became the primary stabilizer in the ACL, taking on 85% of the total force applied to the ACL during anterior drawer. This continued to increase up to 91% in the 6-month boars. In the barrows, the AM bundle contributed 70% and 77% at 4.5 and 6-months, respectively. To compare boar and barrow bundle contributions across age, the difference in AM and PL percent contribution was plotted over time, with negative values indicating greater PL bundle forces and positive values indicating greater AM bundle forces (Fig 6E). There was a statistically significant age effect seen (p=0.012), with AM bundles becoming more dominant across growth for both boars and barrows. Furthermore, all 4.5-month and 6-month differences in percent bundle contribution were statistically greater than 0, indicating higher AM bundle contribution. There was no statistical significance between boars and barrows (p=0.24), yet there was a more gradual shift in barrows towards higher AM bundle contribution, whereas in boars, the contribution of the AM bundle increased sharply between 3 and 4.5 months.

**Figure 6.**
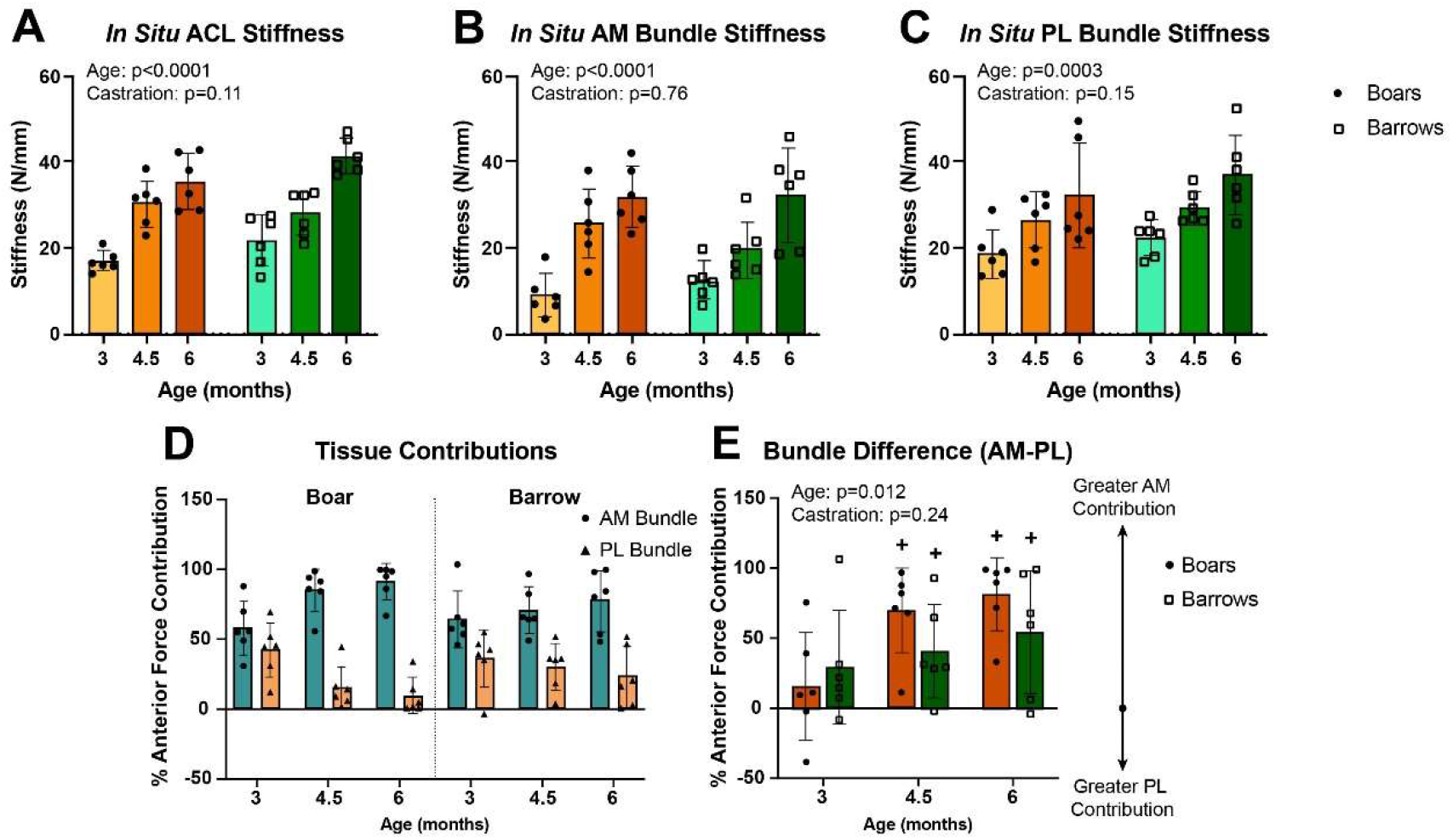
*In situ* ACL, AM bundle, and PL bundle stiffnesses increase throughout skeletal growth and are similar between boars and barrows. (A) ACL, (B) AM bundle, and (C) PL bundle stiffnesses all increase from 3-month to 6-months. ACL and AM bundle stiffnesses calculated during intact kinematics, and PL bundle stiffnesses calculated during the AM bundle deficient kinematic state. (D) Tissue contributions varying across age, with the AM bundle become more dominant in boars and slowly increasing in barrows. (E) Difference in bundle contribution similar in boars and barrows, with both increasing toward AM bundle dominance. Data presented as mean ± 95% confidence interval with main effects from two-way ANOVA shown in graph corner. + indicates p<0.05 relative to zero (indicating AM≠PL) within age groups.

## Discussion

This work explored the influence of neo-natal surgical castration on ACL morphology and function later in youth and early adolescence in a male porcine model. Surgical castration lowers serum testosterone levels by over 95%^21^ and is a standard practice in the swine industry to control for meat quality^35^. The ACL increased considerably in size and stiffness across skeletal growth for both intact males (boars) and castrated males (barrows). Relative to changes in growth, however, there were only subtle differences between the two boars and barrows. Barrows had, on average, smaller overall knee joints along with smaller absolute ACL, AM bundle, and PL bundle volumes, but no difference in ACL or bundle CSA. When accounting for joint size, any differences disappeared. Functionally, the AM bundle percent contribution increased across growth for both groups, yet the boars had a more dramatic shift from equal bundle percent contribution to a large increase in AM bundle percent contribution.

While joint size has not been previously characterized between boars and barrows, it has been shown that barrows have lighter tibias than boars despite similar density and length measurements^36^, indicating greater bone thickness in boars. Differences in body composition between boars and barrows have been well-characterized, with barrows having more fat and less bone and skin weight than boars^36–38^. This finding is similar in humans, where supraphysiological doses of testosterone lead to more fat-free mass and muscle size in men^39^. When looking at changes across skeletal growth, the increase in ACL length, CSA, and volume matches previous findings in both pigs^24,30^ and humans^40–42^. The minor changes in ACL morphology observed in this study, especially after normalizing to the overall joint size are similar to other findings directly relating intact and castrated animals^22^. A previous study measured ACL CSA after 12-weeks in rats castrated as adults and intact male rats and found that the ACL CSA did not differ between the two groups. The volume of the ACL and the AM and PL bundles had more substantial changes between boars and barrows, with barrows having smaller ACLs, AM bundles, and PL bundles than boars. The more substantial changes in ACL, AM bundle, and PL bundle volume could simply be a result of magnifying smaller changes seen in the CSA and length or could be of a result of larger differences near the entheses of the ACL. However, normalizing to joint size also diminished these differences in volume, indicating that surgical castration may influence absolute size, but may not influence normalized joint properties that can be used to compare individuals.

Regarding mechanics, both boars and barrows had increasing ACL, AM bundle, and PL bundle stiffnesses across skeletal growth, yet no significant differences were observed between boars and barrows. While the stiffness of the joint and the tissues increased throughout skeletal growth, there were also minor increases across growth for boars and barrows in absolute joint laxity as measured by APTT, but similar laxity values once normalized to joint size. Furthermore, the barrows had on average, less lax joints and slack, especially at 3 months, yet this disappeared after normalizing to joint size. The normalized laxity values for the boars and barrows only varied 5.8% in the boars and 5.1% in the barrows across ages. Human studies often report raw values of APTT at static loads across ages, such as one study that showed that skeletally mature men have a decreased knee laxity compared to immature men where female knee laxity does not change^43^. This differs in this study where scaled loads were applied and laxity values were similar within boars and barrows. If static loads were applied, then there might have been similar decreases in absolute laxity across skeletal growth.

Previously, the pig model has been used to detect differences in bundle function across skeletal growth and found that the AM bundle takes on more of a dominant load (92% by 6-months in males and 76% by 6-months in females) over the PL bundle^30^. The AM bundle contribution in the barrows matches the 76% AM bundle contribution in female pigs at 6-months yet is higher than boars or females at 3 months (11% and 44%, respectively). Temporally, the female AM bundle contribution increases from 3- to 4.5- to 6-months, boars have a rapid rise between 3- and 4.5-months then plateau, and barrows have higher AM bundle contribution from 3-months and gradually rise over time. Because there are no significant differences in the normalized morphological or stiffness values of the ACL or its bundles, it is interesting that there is a slight, yet not statistically significant, difference in the bundle contribution to anterior loading, where the bundles in the barrow behave more similar to female pigs than to intact male pigs (Figure S3). When looking at the bundle function across skeletal growth and overall joint size, the timing seems to support an earlier maturation in boars than in barrows, yet CSA and stiffness values seems to indicate no difference in ACL maturation. Perhaps testosterone has greater influence on the microstructural properties of the collagen fibers in the ACL, as shown by a decrease in collagen content and increase in collagen fiber diameter in castrated male rabbits^44^. Given that the rise of testosterone in boars starts around 4-months^45^, the slight difference in timing of functional changes could be influenced more by testosterone especially between the 3-month and 4.5-month range. Additional variables could also be at play when making comparisons between gilts, boars, and barrows, such as fluctuations in other female sex hormones^14–18^.

Results from this work suggest that surgical castration at an early age has minor impacts on the ACL structure and function several months later into early adolescence. The current results cannot explain the continued rise in ACL, AM bundle, and PL bundle CSA in intact male pigs in early adolescence compared to a plateau in ACL and PL bundle CSA in female pigs over the same age period. Other factors exist, such as the equilibrium of the conversion between androgens and estrogens via the aromatase enzyme^46^. With this conversion, castration can also decrease serum estrogens in males. With other studies showing that females have higher joint laxity than males, which is a known risk factor for ACL injury^47,48^, the previous work in this pig model showing a 38% greater APTT for female pigs at 18-months^30^, and this work showing similar normalized laxity values for boars and barrows, the future work should focus on the need to directly measure multiple hormonal factors in females.

There are some limitations to the present study that may provide new pathways to understanding the role of testosterone in the developing male ACL. The primary limitation is that while the pig is a good model for understanding human knee function due to its similar anatomical structures and size, no animal model completely replicates the human condition. Another limitation of this study is the loss of stabilizing muscles surrounding the knee during applied loads. Testosterone has been shown to have more drastic impacts on muscle mass and strength^49,50^, so there still could be a greater impact of testosterone on *in vivo* joint stability between boars and barrows due to greater muscle stabilization. When looking at the functional characteristics of the ACL and its bundles, only sub-failure properties were assessed. A previous study in rats discovered that castrated rats had lower load-to-failure forces and ultimate tensile stresses than intact males, yet these failure properties were not assessed in this current study^22^. Furthermore, there was a high variability in the barrow bundle percent contributions. Due to limitations of specimen collection, there was also a lack of serum testosterone measurements in the boars, so associative analyses could not be done. Longer-term data in older animals would perhaps show a long-term impact of testosterone after its normal rise at 4 months. Sex hormones are generally complex in nature, with independent hormones rarely impacting macroscopic properties, and studying the interaction between fluctuating levels of serum sex hormones will prove useful in attempting to understand sex-specific differences throughout skeletal growth and ACL injury risk.

In conclusion, surgical castration had subtle impacts on joint and ACL morphology and function across skeletal growth. Across skeletal growth, the boars and barrows had increasing joint sizes, ACL CSA values and volumes, and stiffnesses. Between boars and barrows, the joint size, ACL, AM bundle, and PL bundle volumes were smaller in the barrow group than the boars across ages, but not the ACL CSA, stiffness, or bundle contribution. The barrows had a more gradual shift in bundle function toward AM-dominance in resisting load than boars, although this was not statistically significant due to high variability in the barrows. Thus, these results show that early and sustained loss in testosterone leads to some differences in ACL morphology but may not influence measures that may indicate increased injury risk, such as CSA or force in the bundles in response to applied loads. This work is beneficial in understanding how hormones might impact ACL performance and injury risk throughout male development.

## Acknowledgments

We would like to thank the Swine Education Unit at North Carolina State University, Laboratory Animal Resources at North Carolina State University, and the Biomedical Research Imaging Center at the University of North Carolina – Chapel Hill for their contributions to this work. Research in the publication was supported by the National Institute of Arthritis and Musculoskeletal and Skin Diseases of the US National Institutes of Health (R01 AR071985 and F31AR077997). The content is solely the responsibility of the authors and does not necessarily represent the official views of the National Institutes of Health. Additional funding for this study was provide in part by the National Science Foundation (DGE-1746939).

