## Supplemental Material for "Neo-natal castration leads to subtle differences in porcine anterior cruciate ligament morphology and function in adolescence"

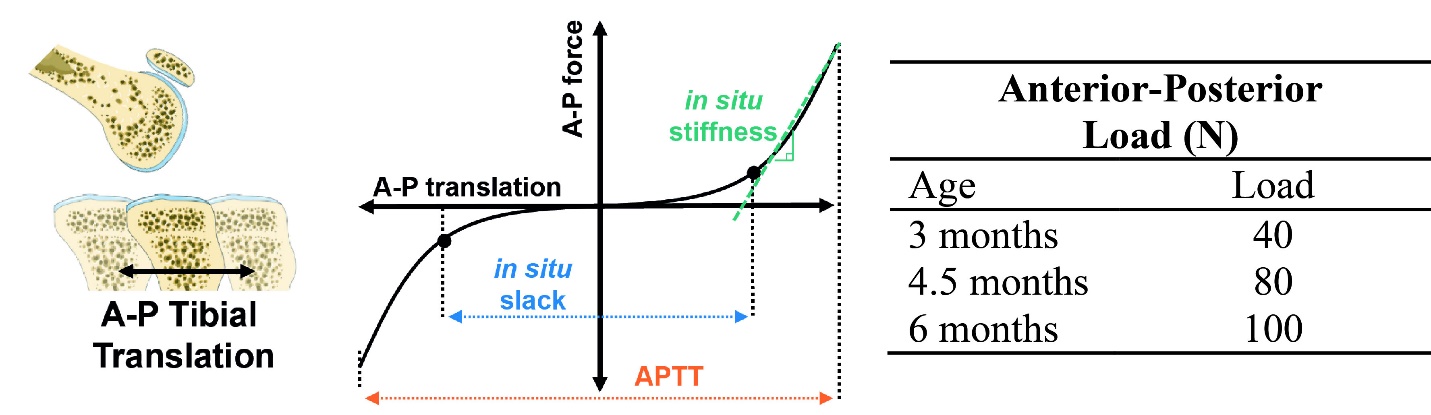

**Figure S1.** AP scaled loading and schematic representation of calculated joint biomechanics measurements, including *in situ* stiffness, *in situ* slack, and APTT.

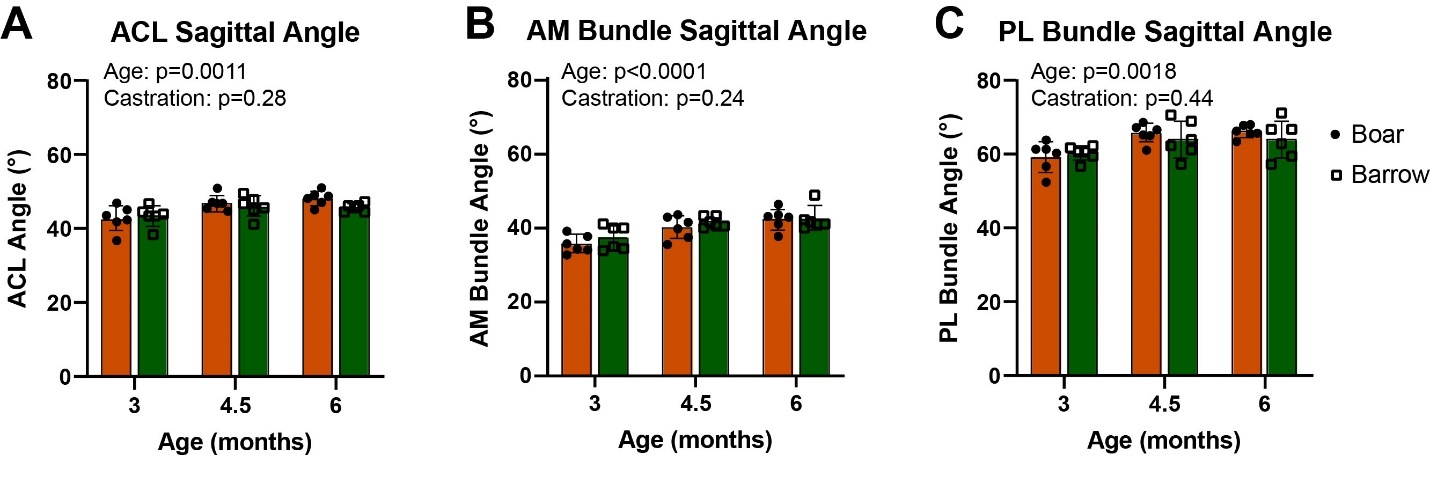

**Figure S2.** Boar and barrow ACL and bundle sagittal orientation increased throughout skeletal growth and adolescence in similar ways. The sagittal angle of the (A) ACL, (B) AM bundle, and (C) PL bundle increased steadily across adolescence, and increases were similar between boars and barrows. Data presented as mean ± 95% confidence interval with main effects from two-way ANOVA shown in graph corner.

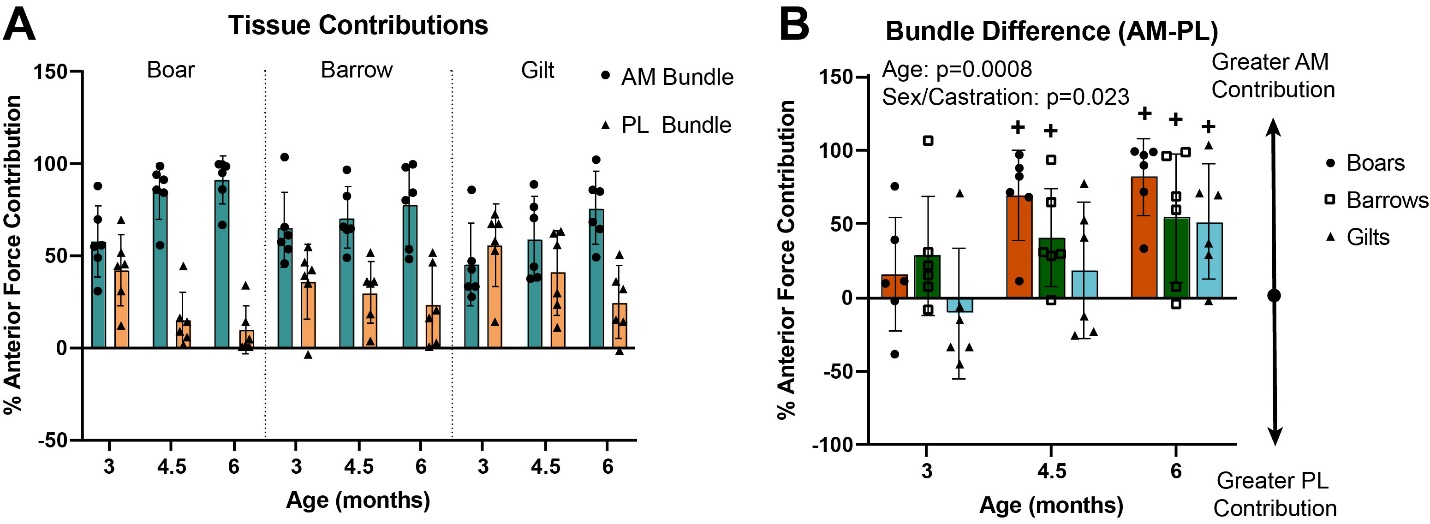

**Figure S3.** Male (boar and barrow) and female (gilt) pig bundle contribution change at different rates. (A) Tissue contributions varying across age, with the AM bundle become more dominant in boars and slowly increasing in barrows and gilts. (B) Difference in bundle contribution similar in boars and barrows, with both increasing toward AM bundle dominance. Data presented as mean ± 95% confidence interval with main effects from two-way ANOVA shown in graph corner. + indicates p<0.05 relative to zero (indicating AM≠PL) within age groups.

**Table S1.** Joint size measurements for boars and barrows at all ages.

|  | **Bicondylar Width (mm)** | | **Tibial Plateau CSA (mm^2^)** | | **Anterior-Posterior Width (mm)** | |
| --- | --- | --- | --- | --- | --- | --- |
| Age | Boar | Barrow | Boar | Barrow | Boar | Barrow |
| 3 months | 54 ± 2 | 50 ± 1 | 3252 ± 252 | 2829 ± 132 | 47 ± 2 | 43 ± 2 |
| 4.5 months | 58 ± 3 | 57 ± 4 | 4028 ± 353 | 3898 ± 357 | 53 ± 2 | 50 ± 2 |
| 6 months | 65 ± 3 | 60 ± 4 | 5328 ± 319 | 4612 ± 499 | 60 ± 2 | 56 ± 2 |

Data presented as mean ± standard deviation.

**Table S2.** ACL, AM bundle, and PL bundle CSA values for boars and barrows at all ages.

|  | **ACL CSA (mm^2^)** | | **AM Bundle CSA (mm^2^)** | | **PL Bundle CSA (mm^2^)** | |
| --- | --- | --- | --- | --- | --- | --- |
| Age | Boar | Barrow | Boar | Barrow | Boar | Barrow |
| 3 months | 30 ± 3 | 31 ± 6 | 11 ± 2 | 13 ± 4 | 15 ± 4 | 17 ± 4 |
| 4.5 months | 45 ± 4 | 40 ± 6 | 21 ± 4 | 17 ± 4 | 19 ± 3 | 20 ± 4 |
| 6 months | 63 ± 9 | 56 ± 10 | 30 ± 5 | 25 ± 7 | 29 ± 5 | 28 ± 5 |

Data presented as mean ± standard deviation.

**Table S3.** ACL, AM bundle, and PL bundle length values for boars and barrows at all ages.

|  | **ACL Length (mm)** | | **AM Bundle Length (mm)** | | **PL Bundle Length (mm)** | |
| --- | --- | --- | --- | --- | --- | --- |
| Age | Boar | Barrow | Boar | Barrow | Boar | Barrow |
| 3 months | 27 ± 2 | 24 ± 1 | 32 ± 3 | 30 ± 1 | 23 ± 1 | 19 ± 1 |
| 4.5 months | 29 ± 2 | 28 ± 2 | 34 ± 2 | 33 ± 2 | 24 ± 1 | 22 ± 2 |
| 6 months | 33 ± 3 | 31 ± 1 | 38 ± 3 | 36 ± 2 | 25 ± 1 | 24 ± 1 |

Data presented as mean ± standard deviation.

**Table S4.** ACL, AM bundle, and PL bundle volumes for boars and barrows at all ages.

|  | **ACL Volume (mm^3^)** | | **AM Bundle Volume (mm^3^)** | | **PL Bundle Volume (mm^3^)** | |
| --- | --- | --- | --- | --- | --- | --- |
| Age | Boar | Barrow | Boar | Barrow | Boar | Barrow |
| 3 months | 864 ±1 24 | 788 ± 169 | 390 ± 53 | 418 ± 104 | 454 ± 99 | 412 ± 91 |
| 4.5 months | 1420 ± 218 | 1228 ± 158 | 829 ± 184 | 636 ± 118 | 547 ± 54 | 532 ± 48 |
| 6 months | 2317 ± 333 | 1869 ± 258 | 1341 ± 186 | 1005 ± 240 | 866 ± 222 | 907 ± 187 |

Data presented as mean ± standard deviation.

**Table S5.** *In situ* joint stiffness, slack, and normalized *in situ* joint slack for boars and barrows at all ages.

|  | ***In situ* Joint Stiffness (N/mm)** | | ***In situ* Joint Slack (mm)** | | **Normalized *In situ* Joint Slack (mm)** | |
| --- | --- | --- | --- | --- | --- | --- |
| Age | Boar | Barrow | Boar | Barrow | Boar | Barrow |
| 3 months | 15.6 ± 3.7 | 20.3 ± 6.3 | 5.5 ± 0.8 | 4.2 ± 1.0 | 0.12 ± 0.01 | 0.10 ± 0.02 |
| 4.5 months | 29.1 ± 5.8 | 27.9 ± 4.9 | 6.1 ± 0.8 | 5.3 ± 1.1 | 0.12 ± 0.02 | 0.11 ± 0.02 |
| 6 months | 33.8 ± 6.1 | 38.8 ± 2.7 | 5.9 ± 0.8 | 5.6 ± 0.7 | 0.10 ± 0.01 | 0.10 ± 0.01 |

Data presented as mean ± standard deviation.

**Table S6.** Raw and normalized joint AP tibial translation for boars and barrows at all ages.

|  | **APTT (mm)** | | **Normalized APTT (mm)** | |
| --- | --- | --- | --- | --- |
| Age | Boar | Barrow | Boar | Barrow |
| 3 months | 9.5 ± 0.7 | 7.7 ± 1.3 | 0.20 ± 0.02 | 0.18 ± 0.03 |
| 4.5 months | 10.5 ± 1.1 | 9.8 ± 1.6 | 0.20 ± 0.03 | 0.20 ± 0.03 |
| 6 months | 10.7 ± 0.9 | 10.0 ± 0.7 | 0.18 ± 0.02 | 0.18 ± 0.02 |

Data presented as mean ± standard deviation.

**Table S7.** ACL, AM bundle, and PL bundle *in situ* stiffnesses for boars and barrows at all ages.

|  | ***In situ* ACL Stiffness (N/mm)** | | ***In situ* AM Bundle Stiffness (N/mm)** | | ***In situ* PL Bundle Stiffness (N/mm)** | |
| --- | --- | --- | --- | --- | --- | --- |
| Age | Boar | Barrow | Boar | Barrow | Boar | Barrow |
| 3 months | 16.8 ± 2.4 | 21.7 ± 5.9 | 8.9 ± 4.9 | 12.3 ± 4.4 | 18.7 ± 5.6 | 22.2 ± 4.1 |
| 4.5 months | 30.4 ± 5.4 | 28.0 ± 5.1 | 25.6 ± 8.2 | 19.6 ± 6.7 | 26.3 ± 6.5 | 29.3 ± 3.8 |
| 6 months | 35.5 ± 6.5 | 41.4 ± 4.1 | 31.7 ± 7.2 | 32.1 ± 11.1 | 32.2 ± 12.1 | 37.1 ± 9.3 |

Data presented as mean ± standard deviation.

**Table S8.** AM bundle and PL bundle percent contribution to total ACL anterior force during anterior loading.

|  | **AM Bundle Contribution**  **(% of ACL Anterior Force)** | | **PL Bundle Contribution**  **(% of ACL Anterior Force)** | |
| --- | --- | --- | --- | --- |
| Age | Boar | Barrow | Boar | Barrow |
| 3 months | 57.9 ± 19.3 | 64.4 ± 20.2 | 42.1 ± 19.3 | 35.6 ± 20.2 |
| 4.5 months | 84.8 ± 15.3 | 70.3 ± 16.6 | 15.2 ± 15.3 | 29.7 ± 16.6 |
| 6 months | 90.8 ± 12.9 | 77.1 ± 21.9 | 9.2 ± 12.9 | 22.9 ± 21.9 |

Data presented as mean ± standard deviation.
